# Bringing Elton and Grinnell Together: a quantitative framework to represent the biogeography of ecological interaction networks

**DOI:** 10.1101/055558

**Authors:** Dominique Gravel, Benjamin Baiser, Jennifer A. Dunne, Jens-Peter Kopelke, Neo D. Martinez, Tommi Nyman, Timothée Poisot, Daniel B. Stouffer, Jason M. Tylianakis, Spencer A. Wood, Tomas Rosling

## Abstract

Biogeography has traditionally focused on the spatial distribution and abundance of species. Both are driven by the way species interact with one another, but also by the way these interactions vary across time and space. Here, we call for an integrated approach, adopting the view that community structure is best represented as a network of ecological interactions, and show how it translates to biogeography questions. We propose that the ecological niche should encompass the effect of the environment on species distribution (the Grinnellian dimension of the niche) and on the ecological interactions among them (the Eltonian dimension). Starting from this concept, we develop a quantitative theory to explain turnover of interactions in space and time *i.e*. a novel approach to interaction distribution modelling. We apply this framework to host parasite interactions across Europe and find that two aspects of the environment (temperature and precipitation) exert a strong imprint on species co-occurrence, but not on species interactions. Even where species co-occur, interaction proves to be stochastic rather than deterministic, adding to variation in realized network structure. We also find that a large majority of host-parasite pairs are never found together, thus precluding any inferences regarding their probability to interact. This first attempt to explain variation of network structure at large spatial scales opens new perspectives at the interface of species distribution modelling and community ecology.

## Introduction

Community ecology is *the study of the interactions that determine the distribution and abundance of organisms* (Krebs, 2009). Despite a general consensus on this definition (Scheiner & Willig, 2007), research on variation in community structure has mostly focused on the spatial and temporal turnover of species composition (Anderson *et al.*, 2011), neglecting variation in the way species interact with each other despite accumulating empirical evidence that this is a major source of diversity (Poisot *et al.*, 2015b). Given this omission, it is perhaps not surprising that biogeographers are still struggling to establish whether interactions actually impact the distribution of species at large spatial scales (Wisz *et al.*, 2012; Kissling *et al.*, 2012): treating interactions as fixed events neglects a large part of the complexity of empirical communities, and will most likely deliver underwhelming results. Recent attempts at accounting for interactions in species distribution models (Pollock *et al.*, 2014; Pellissier *et al.*, 2013) have brought some methodological advances, but are not sufficient for two reasons. First, these techniques are still based on a ‘species-based’ approach to communities, where interactions are merely treated as fixed covariates affecting distribution. Second, they failed to provide a conceptual step forward, both in their treatment of interactions and in the quality of the predictions they make.

Network approaches offer a convenient representation of communities because they simultaneously account for species composition and their interactions. Species are represented as nodes, so that networks already encompass all the information used by current approaches; in addition, interactions are represented by links, so that networks provide additional, higher-order information on community structure. To date, studies of network diversity have mostly been concerned with the distribution of interactions within locations, and less so with variation among locations (Dunne, 2006; Bascompte & Jor-dano, 2007; Ings *et al.*, 2009; Kéfi*et al.*, 2012). There is, however, ample evidence that interaction networks vary in space and time (Lalibertä & Tylianakis, 2010; Poisot *et al.*, 2012; Albouy *et al.*, 2014; Poisot *et al.*, 2016b; Trøjelsgaard *et al.*, 2015), even though there is no common framework with which to generalize these results. Metacommunity theory provides explanations for variation in the distribution of the nodes (Gravel *et al.*, 2011; Pillai *et al.*, 2011), but there is no such explanation to the variation of node and link occurrences. Consequently, we urgently need a conceptual framework to formalize these observations, as it is the only way towards fulfilling the goal of community ecology: providing cogent predictions about, and understanding of, the structure of ecological communities.

Given the historically different approaches to modelling the distributions of species vs. interactions, there is a clear need to bring the two together. Here, we offer an integrated approach to do so, adopting the view that community structure is best represented as a network of ecological interactions. Based on this idea, we propose a new description of the basic concept of the ecological niche that integrates the effect of the environment on species distribution and on the ecological interactions among them. Building on this concept, we develop a quantitative theory to explain turnover of interactions in space and time. We first present the conceptual framework, and then formalize it mathematically, using a probabilistic model to represent the sampling of the regional pool of interactions. At the level of species pairs, the statistical approach could be conceived as an interaction distribution model. At the community level, the approach provides a likelihood-based method to compare different hypotheses of network turnover. As an illustrative example, we apply this novel framework to a large data set on host-parasite interactions across Europe and find that two aspects of the environment (temperature and precipitation) exert a strong imprint on species co-occurrence, but not on species interactions. The network structure changes systematically across the latitudinal gradient, with a peak of connectance at intermediate latitudes.

## The two dimensions of community structure

The problem of community assembly is often formulated as *how are species sampled from a regional pool to constitute a local community (Götzenberger et al., 2012)?* This question could be rewritten to address the problem of network assembly, as *how do samples from a regional pool of interactions constitute a local interaction network?* An illustration of this problem for a food web is provided in Fig. 1. The regional pool of interactions, the *metaweb*, represents potential interactions among all species that could be found in a given area. In this particular case, there are 275 nodes, and 1173 links among the plants (52 nodes), herbivores (96 nodes), and parasitoids (127 nodes) from Northern Europe. An instance of a local community is also illustrated, with 45 nodes and 93 interactions. Only 55.0% of all potential interactions (plant-herbivore or herbivore parasitoid combinations) are realized locally, revealing the stochastic nature of ecological interactions. Our objective here is to provide a conceptual framework to explain the sampling of the regional pool of interactions, along with a quantitative method to predict it. The problem could be formalized sequentially by understanding first why only a fraction of the species cooccur locally and second why these species do or do not interact.

**Figure 1.**
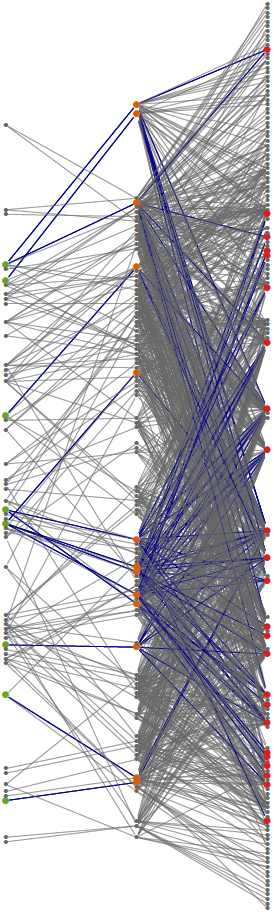
Non-random sampling of the metaweb. Network assembly can be viewed as a sampling process of the regional pool of potential interactions. Species (indicated by colored nodes) are sampled first, and among the species found in the local network, only some interactions (indicated by blue links) occur. We characterize these sampling processes with the quantitative framework proposed in this paper. As a concrete illustration of metaweb sampling, we here show a local interaction network among Salix (left/green), gallers (center/red), and parasitoids (red/blue). The metaweb was constructed by aggregating interactions observed across 370 local networks.

There are multiple causes of spatial turnover of species co-occurrence. The first and most-studied driver is the effect of variation in the abiotic environment on species performance. Combined with specific responses in demography, it generates variation among sites by selecting the locally fittest species (Leibold *et al.*, 2004). Stochasticity plays an additional role, either because colonization and extinction events (Hanski, 1999) are inherently unpredictable or because strong non-linear feedbacks in community dynamics generate alternative transients and equilibria (Chase, 2007; Vellend *et al.*, 2014). Analyses of community turnover are usually performed with data represented in a table with rows corresponding to sites (or measurements) and columns to species. Metrics of beta diversity quantify the variance of this community data (Legendre *et al.*, 2005). Traditional approaches rely on measures of dissimilarity among communities, such as the Jaccard or Bray-Curtis indices. More recent approaches decompose total variation of the community data into species and site contributions to beta diversity (Legendre & De Caceres, 2013), and further partition it into dissimilarity due to changes in species richness and dissimilarity due to actual species turnover (Baselga, 2010). Even though these methods compare whole lists of species among sites or measurements, they remain fundamentally “species-based”, since they report variation within columns. None of them explicitly considers variation of associations (i.e., of pairs or higher-order motifs-Stouffer *et al.* 2007).

Similarly, we are now getting a better understanding of interaction turnover. As mentioned above, in the network approach to community structure, species and interactions are represented by nodes and links, respectively. Associations can also be represented by matrices in which entries represent the occurrence or intensity of interactions among species (rows and columns). Network complexity is then computed as the number of interactions (in the case of binary networks) or interaction diversity (in the case of quantitative networks, Bersier *et al.* 2002). Variability in community structure consequently arises from the turnover of species composition, along with turnover of interactions among pairs of species. The occurrence and intensity of interactions could vary because of the environment, species abundance, and higher-order interactions (Poisot *et al.*, 2015b). Variation in community composition can be independent of variation of ecological interactions, suggesting that species and interaction distribution may well respond to different drivers (Poisot *et al.*, 2012).

The “niche” is by far the dominant concept invoked to explain species distributions and community assembly, from the local to the global scale. Following Hutchinson 1957, the niche is viewed as the set of environmental conditions allowing a population to establish and persist (see also Holt 2009). Community turnover arises as a result of successive replacement of species along an environmental gradient, in agreement with the Gleasonian view of communities (Gleason, 1926). The concept is straightforward to put into practice with species distribution models, as it maps naturally on available distributional and environmental data. Consequently, a vast array of statistical tools have been developed to implement it (e.g. BIOMOD Thuiller 2003, MaxEnt Phillips *et al.* 2006). It is however much harder to account for ecological interactions within this approach (Townsend *et al.*, 2011). As such, these interactions are often viewed as externalities constraining or expanding the range of environmental conditions required for a species to maintain a viable population (Pulliam, 2000; Soberón, 2007).

Interestingly, the ecological network literature also has its own “niche model” to position a species in a community (Williams & Martinez, 2000). The niche of a species in this context represents the multidimensional space of all of its interactions. Each species is characterized by a niche position, an optimum and a range over three to five different niche axes (Williams & Martinez, 2000; Eklöf *et al.*, 2013). The niche model of food web structure and its variants have successfully explained the complexity of a variety of networks, from food webs to plant-pollinator systems (Allesina *et al.*, 2008; Williams *et al.*, 2010; Eklöf *et al.*, 2013). This conceptual framework is, however, limited to local communities, and does not provide any explanation for the turnover of network structure along environmental gradients.

## The integrated niche

Despite several attempts to update the concept of the ecological niche, ecologists have not moved far beyond the “n-dimensional hypervolume” defined by Hutchinson. Despite its intuitive interpretation and easy translation into species distribution models (Boulangeat *et al.*, 2012; Blonder *et al.*, 2014), the concept has been frequently criticized (Hardin, 1960; Peters, 1991; Silvertown, 2004), and several attempts have been made to expand and improve it (Pulliam, 2000; Chase & Leibold, 2003; Soberón, 2007; Holt, 2009; McInerny & Etienne, 2012b).

Part of the problem surrounding the niche concept has been clarified with the distinction between Eltonian and Grinnellian definitions (Chase & Leibold, 2003). The Grinnellian dimension of the niche is the set of environmental conditions required for a species to maintain a population in a location. The Grinnellian niche is intuitive to apply, and constitutes the conceptual backbone of species distribution models. The Eltonian niche, on the other hand, is the effect of a species on its environment. While this aspect of the niche is well known by community ecologists, it is trickier to turn into predictive models. Nonetheless, the development of the niche model of food web structure (Williams & Martinez, 2000) and its parameterization using functional traits (Gravel *et al.*, 2013; Bartomeus *et al.*, 2016) made it more operational.

These perspectives are rather orthogonal to each other, and this has resulted in considerable confusion in the literature (McInerny & Etienne, 2012a). Chase & Leibold 2003 attempted to reconcile with the following definition: *“[The niche is] the joint description of the environmental conditions that allow a species to satisfy its minimum requirements so that the birth rate of a local population is equal to or greater than its death rate along with the set of per capita effects of that species on these environmental conditions”*. Their representation merges zero-net-growth isoclines delimiting the Grinnellian niche (“when does the population persists?”) with impact vectors delimiting the Eltonian niche (“what is the per-capita impact?”). While this representation has been very influential in local-scale community ecology (the resource-ratio theory of coexistence, Tilman 1982), it remains impractical at larger spatial scales because of the difficulties in measuring it. The absence of any mathematical representation of the niche that can be easily fit to ecological data may explain why biogeographers are still struggling to develop species distribution models that also consider ecological interactions. Thus, a more integrative description of the niche will be key to understand spatial and temporal turnover in community structure.

We propose to integrate the two perspectives of the niche using a visual representation of both components (Fig. 2). The underlying rationale is that, in addition to the environmental constraints on demographic performance (Fig.2 top panel), any organism requires resources to meet its metabolic demands and to sustain reproduction (Fig. 2 nodes in network of bottom panel). Abiotic environmental axes are any non-consumable factors affecting the demographic performance of an organism. Alternatively, the resource axes are traits of the resources that allow interactions with the consumer. The niche can therefore be viewed as the set of abiotic environmental variables (the Grinnellian component) along with the set of traits (the Eltonian component) that allow a population to establish and to persist at a location. Accordingly, each species can be characterized by an optimal position along both the environmental (x-axis) and the trait (y-axis) plane. The integrated niche is then the hypervolume where interactions can occur and sustain a population.

**Figure 2.**
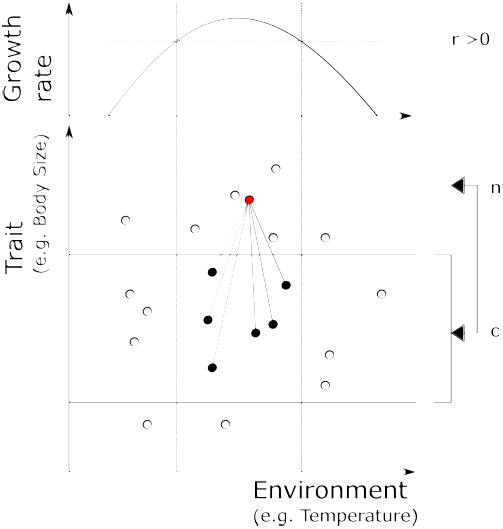
Visual representation of the integrated niche. In biogeography, the niche is considered the set of environmental conditions where the intrinsic growth rate r is positive (Holt, 2009). The horizontal axis represents an environmental gradient impacting the growth of the focal species (in red). The location of each species along this gradient represents their optimum, and the vertical dotted lines represent the limits of the Grinnellian niche of the focal species. In food web ecology, the Eltonian niche represents the location of a species in the food web, as determined by its niche position (n) and its niche optimum (c). The vertical axis represents a niche gradient, for example a trait such as body size. The location of each species along this gradient represents their niche position. The focal species will feed only on prey species occupying niche locations within a given interval around the optimum, represented by the horizontal lines. The integrated Grinnellian and Eltonian niche corresponds to the square in the middle where an interaction is possible owing to a match of traits and spatial distribution. According to our probabilistic framework, the central square represents the area where the joint probability of observing co-occu and interactions is positive.

This approach radically changes the representation of the niche, putting species distributions and ecological interactions into the same formalism. Moreover, it allows the limits of the niche axes to be independent of each other (as in the example in Fig. 2), or to interact. For instance, the optimal prey size for predatory fishes could decline with increasing temperature (Gibert & DeLong, 2014), which would make diet boundaries functions of the environment. Alternatively, we could also consider that the growth rate of the predator changes with the size of its prey items, thereby altering the environmental boundaries.

## A probabilistic representation of interaction networks in space

We now formalize the integrated niche with a probabilistic approach to interactions and distributions. In particular, we seek to represent the probability that an interaction between species *i* and *j* occurs at location *y*. We define *L_ijy_* as a stochastic variable, and focus on the probability that this event occurs, *P*(*L_ijy_*). We note that the occurrence of an interaction is dependent on the co-occurrence of species *i* and *j*. This argument might seem trivial at first, but the explicit consideration of this condition in the probabilistic representation of ecological interactions will prove instrumental to understanding their variation. We define *X*_iy_ as a stochastic variable representing the occurrence of species *i* at location *y*. The quantity we seek to understand is the probability of a joint event, conditional on the set of environmental conditions *E_y_*:

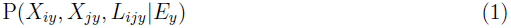

Or simply said, the probability of observing both species *i* and *j* plus an interaction between *i* and *j* given the conditions *E*_*y*_ at location *y*. This probability could be decomposed into two parts using the product rule of probabilities:

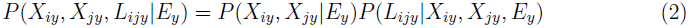

The first term on the right-hand side of the equation is the probability of observing the two species co-occurring at location *y*. It corresponds to the Grinnellian dimension of the niche. The second term represents the probability that an interaction occurs between species *i* and *j*, given that they are co-occurring. This predicate can be refined using information on trait distribution and trait matching rules ((Bartomeus *et al.*, 2016)). Above, we referred to this entity as the “metaweb” and it corresponds to the Eltonian dimension of the niche. Below, we will see how this formalism can be directly fit to empirical data. But before turning to an application, we will discuss the interpretation of different variants of these two terms.

### Variants of co-occurrence

There are several variants to the co-occurrence probability, representing different hypotheses concerning temporal and spatial variation in network structure (see the explicit formulations in Table 1). The simplest model relates the probability of co-occurrence directly to the environment, *P*(*X_iy_*, *X_jy_*|*E*_*y*_). In this situation, there are no underlying assumptions about the ecological processes responsible for co-occurrence. It could arise because interactions constrain distribution (Pollock *et al.*, 2014; Cazelles *et al.*, 2016) or, alternatively, because of environmental requirements shared between *i* and *j*. In the former case, species are not independent of each other and the conditional occurrence must be accounted for explicitly, *P*(*X*_*iy*_, *X*_*jy*_|*E*_*y*_) = *P*(*X_iy_*|*E_y_*, *X_jy_*)*P*(*X*_*jy*_|*E_y_*). In the latter case, species are independent, and only the marginal occurrence must be accounted for, *P*(*X_ijy_*|*E_y_*) = *P*(*X_iy_*|*E_y_*)*P*(*X_jy_*|*E_y_*).

**Table 1:** Summary of model comparison for the interaction between the leaf folder Phyllocolpa prussica) and the parasitoid Chrysocharis elongata.

| # | Metaweb model | Co-occurrence model | LL | npars | AIC |
| --- | --- | --- | --- | --- | --- |
| 1 | $P(L_{ijy})$ | $P(X_{iy}, X_{jy}\vert E_y)$ | -71.1 | 6 | 154.2 |
| 2 | $P(L_{ijy}\vert X_{iy}, X_{jy})$ | $P(X_{iy}, X_{jy}\vert E_y)$ | -65.7 | 6 | 143.4 |
| 3 | $P(L_{ijy}\vert X_{iy}, X_{jy}, E_y)$ | $P(X_{iy}, X_{jy}\vert E_y)$ | -65.6 | 10 | 151.3 |
| 4 | $P(L_{ijy}\vert X_{iy}, X_{jy}, E_y)$ | $P(X_{iy}, X_{jy})$ | -84.5 | 6 | 183 |
| 5 | $P(L_{ijy}\vert X_{iy}, X_{jy}, E_y)$ | $P(X_{iy})P(X_{jy})$ | -80.7 | 7 | 173.4 |
| 6 | $P(L_{ijy}\vert X_{iy}, X_{jy}, E_y)$ | $P(X_{iy}\vert E_y)P(X_{jy}\vert E_y)$ | -68.8 | 15 | 167.6 |

The co-occurrence probability itself could depend on ecological interactions. This should be viewed as the realized component of the niche (i.e. the distribution when accounting for species interactions). Direct pairwise interactions such as competition, facilitation, and predation have long been studied for their impact on co-distribution (e.g. Diamond 1975; Connor & Simberloff 1979; Gotelli 2000). Second-and higher-order interactions (e.g. trophic cascades) could also affect co-occurrence. Co-occurrence of multiple species embedded in ecological networks is a topic of its own, however, and is influenced by both network topology and species richness (Cazelles *et al.*, 2016). Not only direct interactions influence co-occurrence, but indirect interactions do as well (e.g. plant species sharing an herbivore, or herbivores sharing parasitoids, could repel each other in space Holt & Lawton 1993). The impact of direct interactions and first-order indirect interactions on co-occurrence tends to vanish with increasing species richness in the community. Further, co-occurrence is also influenced by the covariance of interacting species to an environmental gradient (Cazelles *et al.*, 2015). Because of the complexity of relating co-occurrence to the structure of interaction networks, we will focus here on the variation of interactions and not on their distribution, and leave this specific issue for the Perspectives section and future research.

### Variants of the metaweb

There are also variants of the metaweb. First, most documented metawebs have thus far considered ecological interactions to be deterministic, rather than probabilistic (e.g. Havens 1992; Wood *et al.* 2015). Species are assumed to interact whenever they are found together in a location, independent of their local abundance and the local environment. In other words, *P*(*L_ijy_*|*X*_*ijy*_ = 1) = 1 and *P*(*L_ijy_*|*X_ijy_* = 0) = 0. This approach might be a reasonable approximation if the spatial or temporal scale of sampling and inference is so large that the probability of observing at least one interaction converges to unity. In this scenario, network variation arises solely from species distributions.

Second, ecological interactions could also vary with the environment, so that *P*(*L_ijy_* |*E_y_*). Although it is rare to see a conditional representation of pairwise ecological interactions, experimental studies have frequently revealed interactions to be sensitive to the environment. For instance, (McKinnon *et al.*, 2010) showed that predation risks of shorebirds vary at the continental scale, decreasing from the south to the north. It is also common to see increasing top-down control with temperature (e.g. Shurin *et al.* 2012; Gray *et al.* 2015). Effects of the environment on interactions also propagate up the community and influence network structure (Tylianakis *et al.*, 2007; Woodward *et al.*, 2010; Petchey *et al.*,2010).

## Application: continental-scale variation of host-parasite community structure

We now turn to an illustration of our framework with the analysis of an empirical dataset of host-parasite networks sampled throughout the south-north environmental gradient in continental Europe. The focal system consists of local food webs of willows (genus Salix), their galling insects, and the natural enemies (parasitoids and inquilines) of these gallers. Targeting this system, we ask: i) how much does network structure vary across the gradient, and ii) what is the primary driver of network turnover across the gradient?

### Data

Communities of willows, gallers, and parasitoids are species-rich and widely distributed, with pronounced variation in community composition across space. The genus Salix includes over 400 species, most of which are shrubs or small trees (Argus, 1997), and is common in most habitats across the Northern Hemisphere (Skvortsov, 1999). Willows support a highly diverse community of herbivorous insects, with one of the main herbivore groups being gall-inducing sawflies (Hymenoptera: Tenthredinidae: Nematinae: Euurina (Kopelke, 1999)). Gall formation is induced by sawfly females during oviposi-tion, and includes marked manipulation of host-plant chemistry by the galler (Nyman & Julkunen-Tiitto, 2000). The enemy community of the gallers includes nearly 100 species belonging to 17 insect families of four orders (Kopelke, 2003). These encompass two main types: inquiline larvae (Coleoptera, Lepidoptera, Diptera, and Hymenoptera) feed primarily on gall tissue, but typically kill the galler larva in the process, while parasitoid larvae (representing many families in Hymenoptera) kill the galler larvae by direct feeding (Kopelke, 2003). In terms of associations between the trophic levels, phylogeny-based comparative studies have demonstrated that galls represent “extended phenotypes” of the gallers, meaning that gall form, location, and chemistry is determined mainly by the galling insects and not by their host plants (Nyman & Julkunen-Tiitto, 2000). Because galler parasitoids have to penetrate a protective wall of modified plant tissue in order to gain access to their victims, gall morphology has been inferred to strongly affect the associations between parasitoids and hosts (Nyman *et al.*, 2007). Thus, the set of parasitoids attacking each host is presumably constrained by the form, size, and thickness of its gall.

Local realizations of the willow-galler-parasitoid network were reconstructed from community samples collected between 1982 and 2010. During this period, willow galls were collected at 370 sites across Central and Northern Europe. Sampling was conducted in the summer months of June and/or July, i.e., during the later stages of larval development. Galler species were identified on the basis of willow host species and gall morphology, as these are distinct for each sawfly species. At each site, galls were randomly collected from numerous willow individuals in an area of about 0.1–0.3 *km*^2^. Some sites were visited more than once, with a total of 641 site visits across the 370 sites. GPS coordinates were recorded for each location; for our analyses, current annual mean temperature and precipitation were obtained from WorldClim using the R package raster (Hijmans, 2015). While other covariates could have also been considered, these two variables are likely representative of the most important axes of the European climate, and are also more easily interpretable than reduced variables obtained, for example, by principal component analysis.

The methods used for rearing parasitoids from the galls have been previously described by Kopelke 2003. In brief, galls were opened to score the presence of galler or parasitoid/inquiline larvae. Parasitoid larvae were classified to preliminary morphospecies, and the identity of each morphospecies was determined by connecting them to adults emerging after hibernation. The galls were reared by storing single galls in small glass tubes (Kopelke, 1985). Hibernation of galls containing parasitoids took place either within the glass tubes or between blotting paper in flowerpots filled with clay granulate or a mixture of peat dust and sand. These pots were stored over the winter in a roof garden and/or in a climatic chamber. In most cases, the matching of larval morphospecies with adult individuals emerging from the rearings allowed the identification of the parasitoids to the species level. Nonetheless, in some cases, individuals could only be identified to one of the (super)families Braconidae, Ichneumonidae, and Chalcidoidea. This was particularly the case when only remains of faeces, vacant cocoons of parasitoids, and/or dead host larvae were found, as was the case when parasitoids had already emerged from the gall. As a result, the largest taxon in the data set, “Chalcidoidea indeterminate”, represents a superfamily of very small parasitoids that are hard to distinguish.

In total, 146,622 galls from 52 Salix taxa were collected for dissection and rearing. These galls represented 96 galler species, and yielded 42,133 individually-identified parasitoids. Of these, 25,170 (60%) could be identified to the species level. Overall, 127 parasitoid and inquiline taxa were distinguished in the material. Data on host associations within subsets of this material have been previously reported by (Kopelke, 1999) and (Nyman *et al.*, 2007). The current study represents the first analysis of the full data set from a spatial perspective.

### Analysis

Computing the probability of observing an interaction involves fitting a set of binomial models and collecting their estimated probabilities. For the sake of illustration, we considered second-order generalized linear models-although more flexible fitting algorithms (e.g. GAM or Random Forest) could equally well be used, as long as the algorithm can estimate the probability for each observation. The data consist of a simple (albeit large and full of zeros) table with the observation of each species, *X_iy_* and *X_jy_*, their co-occurrence, *X_ijy_*, the observation of an interaction *L_ijy_*, and environmental co-variates *E_y_*. Thus, there is one row per pair of species per site. We considered that an absence of a record of an interaction between co-occurring species at a site means a true absence (see below for a discussion on this issue).

We compared three models for the co-occurrence probability. The first one directly models the co-occurrence probability conditional on the local environment, *P*(*X_iy_*,*X_jy_*|*E_y_*) (models are listed at Table 1 and 2). Hence, this model makes no assumptions about the mechanisms driving co-occurrence for any given environment, and instead uses the information directly available in the data. It thereby indirectly accounts for the effect of interactions on co-occurrence, if there are any. The second model considers independent occurrence of species. In this case, we independently fit *P*(*X_iy_*|*E_y_*) and *P*(*X_jy_*|*E_y_*), and we then take their product to derive the probability of co-occurrence. This model should be viewed as a null hypothesis with respect to the first model, since a comparison between the respective models will reveal if there is significant spatial association of the two species beyond a joint response to the shared environment (Cazelles *et al.*, 2016). Finally, the third model assumes that the probability of co-occurrence is independent of the environment and thus constant throughout the landscape. In other words, *P*(*X_iy_*,*X_jy_*) is obtained by simply counting the number of observed co-occurrences divided by the total number of observations. Thus, the comparison between the first and third model allows us to test the hypothesis that co-occurrence is conditional on the environment. Whenever the environment was included as a covariate in the GLM, we considered a second-order polynomial response for both temperature and precipitation in order to account for optima in environmental conditions. There are consequently five parameters for the first model when fitting a given pair of species, 10 parameters for the second, and only one for the third model.

Following the same logic, we compared three models of the interaction probability. The first model conditions the interaction probability on the local environmental variables, *P*(*L_ijy_*|*X_iy_*, *X_jy_*, *E_y_*). Consequently, the model was fit to the subset of the data where the two species co-occur. The second model fits the interaction probability independently of the local environmental variables, *P*(*L_ijy_*|*X_iy_*, *X_jy_*). It corresponds to the number of times the two species were observed to interact when co-occurring, divided by the number of times that they co-occurred. The third model is an extreme case performed only to test the hypothesis that if two species are found to interact at least once, then they should interact whenever they co-occur, *P*(*L_ijy_*|*X_iy_*,*X_jy_*) = 1. While not necessarily realistic, this model tests an assumption commonly invoked in the representation of local networks from the knowledge of a deterministic metaweb. There are consequently five parameters for the first model, a single parameter for the second model and no parameter to evaluate for the third model (where the interaction probability is fixed by the hypothesis).

We fit the different models to each pair of species and recorded the predicted probabilities. The joint probability *P*(*L_ijy_*, *X_iy_*, *X_jy_*) was then computed from Eq. 2, and the likelihood of each observation was computed as *L*(θ*_y_*|*D_ijy_*) = *P*(*L_ij_*, *X_iy_*, *X_jy_*) if an interaction was observed, and as *L*(θ*_ijy_*|*D_ijy_*) = 1 — *P*(*L_ijy_*,*X_iy_*,*X_jy_*) if no interaction was observed. The log-likelihood was summed over the entire dataset to compare the different models by AIC. Not surprisingly, there was a very large number of species pairs for which this model could not be computed, as they simply never co-occurred. For these pairs, we have no information of the interaction probability, and they were consequently removed from the analysis. The log-likelihood reported across the entire dataset was summed over all pairs of species observed to co-occur at least once. Interactions between the first (Salix) and second (gallers) trophic layers and those between the second and third (parasitoids) were considered separately. Finally, we used the full model (in which both co-occurrence and the interaction are conditional on the environment) to interpolate species distributions and interaction probabilities across the entire European continent. We reconstructed the expected network for each location in a 1 X 1 km grid and computed the probabilistic connectance following (Poisot *et al.*, 2016a).

All of the data are openly available in the database *mangal* (Poisot *et al.*, 2015a) and all R scripts for running the analysis, are provided in the Supplementary Material.

### Results

Despite the extensive sampling, many pairs of species were observed to co-occur only a few times. This made it difficult to evaluate interaction probabilities with any reasonable confidence interval. Thus, we start with an example of a single pair of species selected because of its high number of co-occurrences (*N_ij_* = 38): the leaf folder *Phyllocolpa prussica* and the parasidoid *Chrysocharis elongata.* These two fairly abundant species were observed *N*_*i*_ = 49 and *N_j_* = 121 times, respectively, across the 370 sites, and they were found to interact with a marginal probability *P*(*L_ij_*) = 0.55, which means they interacted at 21 different locations. Here, a comparison of model fit (Table 1) reveals that conditioning the interaction probability on local environmental conditions adds no explanatory power beyond a model assuming the same probability of interaction anywhere in space (Model 1 vs Model 2). Moreover, when the two species co-occur, the occurrence of the interaction was insensitive to the environment (Model 2 vs Model 3). Alternatively, climatic variables significantly impacted co-occurrence (Model 3 vs Model 4). The neutral model performed worse than the non-random co-occurrence model (Model 3 vs Model 6). The full model revealed that the greatest interaction probability occurred at intermediate temperature and precipitation, simply because this is where the two species most frequently co-occur (Fig. 3). The probabilities of co-occurrence and interaction can be represented in space, where we found that the highest interaction probability occurred in Central Europe (Fig. 4).

**Figure 3.**
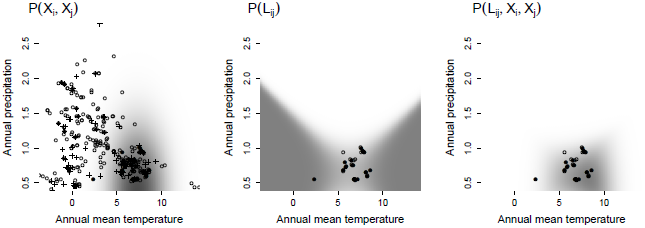
**Probabilistic representation of the interaction probability between a leaf folder(*Phyllocolpa prussica*) and a parasitoid (*Chrysocharis elongata*) across gradients of annual average temperature and annual precipitation**. The representation is based on predictions from Model 3 (see Table 1). In the left panel, open circles represent the absence of both species, whereas closed circles represent co-occurrence and plus signs the occurrence of only one of the two species. In the other two panels, open circles represent co-occurrence but an absence of interaction and closed circles the occurrence of an interaction.

**Figure 4.**
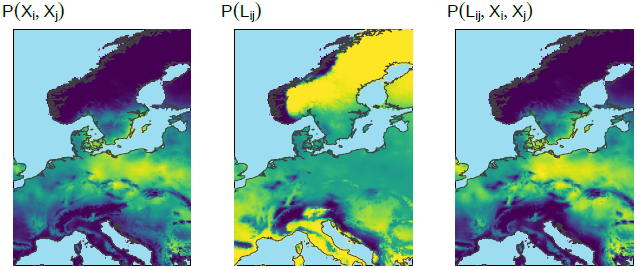
**Probabilistic representation of the interaction probability between a leaf folder (*Phyllocolpa prussica*) and a parasitoid (*Chrysocharis elongata*) across Europe**. The maps are generated from probabilities predicted by the model illustrated inFig. 3

We evaluated each model for all pairs of species in order to better understand the large-scale drivers of network turnover. The results were highly consistent among trophic layers (Salix-gallers and gallers-parasitoids; Table 2). Across all pairs of species, the conditional representation of interactions performed better than the marginal one (Model 1 vs Model 2); that is, interactions did not occur systematically whenever the two species were found co-occurring. Hence, in addition to species turnover, the stochastic nature of interactions contributes to network variability. In total, we recorded 1,173 pairs of interactions, only 290 of which occurred more than five times. Out of these 290 interactions, 143 were systematically detected whenever the two species co-occurred. In the instances when species co-occurred, the two environmental variables considered proved relatively poor predictors of their interactions (Model 2 vs Model 3). Not surprisingly, for both types of interactions (Salix-galler and galler-parasitoid), the log-likelihood increased when the environment was considered. However, the extra number of parameters exceeded the gain in log-likelihood and inflated AIC. Therefore, the most parsimonious model excluded the effect of the environment. On the basis of log-likelihood only, co-occurrence was nonneutral for both Salix-galler and galler-parasitoid interactions. Thus, according to AIC, the best model was the one of non-random co-occurrence (Model 3 vs Model 6) for both types of interactions.

**Table 2:**
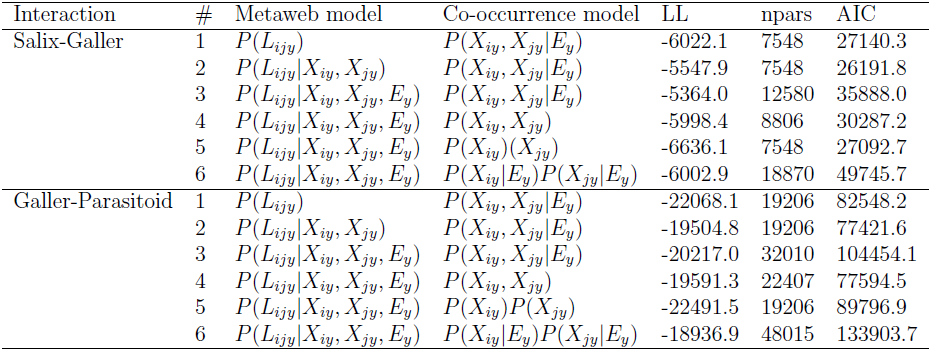
Summary of model comparison for the interaction across all pairs of salix, gallers and parasitoids.

The approach we present not only has implications for understanding the biogeography of pairwise interactions and interaction networks, but also for evaluating the quality of metawebs. We investigated the reliability of the estimated metaweb across the entire dataset wtih summary statistics of species co-occurrence. As mentioned above, across the 17,184 potential pairs of species, only 1,173 pairs interacted in at least a single location, yielding a connectance of 0.068. However, only 4,459 pairs of species were found co-occurring at least once across all locations. There are consequently 12,725 gaps of information in the metaweb (74.1%-see Fig. 5). As we cannot know whether the non-co-occurring species would indeed interact if found together, a more appropriate estimate of connectance would be *C* = 1173/4459 = 0.263. This result reveals that the evaluation of the sampling quality of ecological networks is a problem on its own and well worth further attention.

**Figure 5.**
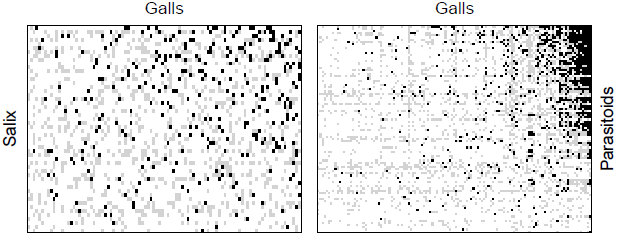
**Representation of the Salix-galler and galler-parasitoid metawebs**. Black cells indicate species pairs for which at least one interaction was recorded, white cells indicate absence of recorded interactions and grey cells show pairs of species never detected at the same site (and hence species pairs for which we have no information on whether they would interact should they co-occur).

Once we had selected the best model based on AIC (Model 3, Table 2), we used it to reconstruct the expected species richness, along with the most likely network for each location. Using this approach, we mapped the expected distribution of network properties across Europe (Fig. 6). For simplicity, we chose to consider connectance as our descriptor of network configuration, as this metric can be easily computed from probabilistic networks (Poisot *et al.*, 2016a) and is also a good proxy for many other network properties (Poisot & Gravel, 2014). Overall, we found a peak in Salix, gallers and parasitoid diversity in Northern Europe. The expected number of interactions roughly followed the distribution of species richness, but accumulated at a rate different from species numbers. Connectance likewise peaked in Northern Europe (Fig. 6).

**Figure 6.**
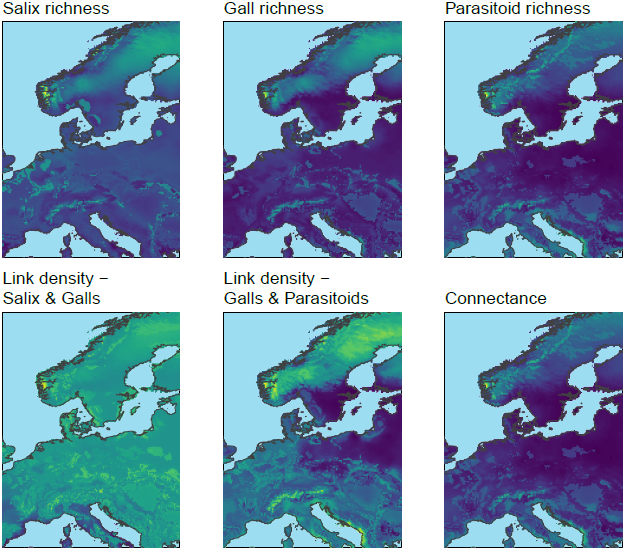
**Mapping the distribution of species richness, the number of links and connectance across Europe.** The representation is based on predictions from Model 3 (see Table 2). Species richness is obtained by summation of individual occurrence probabilities, and link density by summation of interaction probabilities.

## Interpretation

We have proposed that the representation of community structure and its variation in space and time is best captured by the formalism of ecological networks, as both the distribution of species and their interspecific interactions can then be accounted for. We consequently revised the niche concept in order to integrate its abiotic and biotic components that vary over time and space. This integrated niche was represented visually with an ordination of species into an environmental space and a trait space. The fundamental niche of a species is represented as the set of environmental conditions and resources that allow a species to establish in a location, thereby integrating the Eltonian and the Grinnellian components of the niche. We then translated the concept mathematically by investigating the probability of the joint occurrences of species and their interaction, which should be interpreted as an interaction distribution model. We used this approach to characterize the turnover of the structure of ecological interactions in a species-rich tri-trophic network across Western Europe, finding that the primary driver of network variation is the turnover in species composition. To our knowledge, this is the first continental-wide analysis of the drivers of network structure from empirical data on the occurrence of interactions (see Baiser *et al.* 2012; Albouy *et al.* 2014; Poisot *et al.* 2016b).

Applying the framework to our large data set on host-parasite interactions across Europe revealed key features in the interaction between Salix taxa, their herbivores, and the natural enemies of these herbivores. Consistent with a general increase in the diversity of Salix towards boreal areas (Cronk *et al.*, 2015), overall species richness of the networks increased towards the north. The distribution of Salix species richness largely matched those of gallers and parasitoids. These observations within Europe are also matched by the ones found at a global scale for Salix (Argus, 1997; Cronk *et al.*, 2015; Wu *et al.*, 2015) and sawflies (Kouki *et al.*, 1994; Kouki, 1999). Species richness in a common groupd of parasitic wasps, the Ichneumonidae, was originally presumed to show a similar “reversed latitudinal gradient”, but this observation has been recently challenged by findings of rather high ichneumonid diversity in the tropics (Veijalainen *et al.*, 2013). Nevertheless, ichneumonid subfamilies specifically associated with sawflies (Ctenopelmatinae, Tryphoninae) are clearly less diverse in the south.Exactly what processes are responsible for the distribution of species richness at different trophic levels is yet to be established (but see e.g. Roininen *et al.* 2005; Nyman *et al.* 2010; Leppänen *et al.* 2014), but as a net outcome of different latitudinal trends across trophic levels, the distribution of co-occurrence and therefore of potential interactions differed between the first and second layers of feeding links. The correlation between expected Salix and gallers richness was 0.73, while it was 0.58 between gallers and their parasitoids. Therefore, the ratio of herbivores to Salix species is essentially constant across Europe, while each herbivore species is potentially attacked by a and a lower trophic level at the same site was clearly affected by the richer enemy community at higher latitudes. Consequently, overall connectance peaks in Northern Europe (Fig. 6).

In terms of species interacting with each other, our analysis suggests that the environment leaves a detectable imprint on species co-occurrence, but only a slight mark on the occurrence of realized links among species in a specific place: the probability of finding a given combination of species at a higher and a lower trophic level at the same site was clearly affected by the environment, whereas the probability of observing an interaction between the two was not detectably so. This applies to the example species *Phyllocolpa prussica* and *Chrysocharis elongata* (Figs 2 and 3), but also to all species pairs more generally. For the example species pair, the full model revealed that the interaction probability peaks at intermediate temperature and precipitation, simply because this is where the two species co-occur most often. This does not imply that species will always interact when they meet-although this is a basic assumption in most documented metawebs to date (e.g. Havens 1992; Wood *et al.* 2015). Rather, an interaction is a stochastic process whose probability is also influenced by the probability with which species co-occur. What we cannot reliably know is how this stochasticity splits into two sampling processes-i.e., the extent to which a species at the higher trophic level runs into a species at the lower level co-occurring at the site, and the extent to which this interaction is detected by an observer collecting a finite sample. Future work will be required to document the relative importance of these two sources of uncertainty in the occurrence of interactions.

## Perspectives

Evidence that the structure of ecological networks does vary across habitats (e.g. Tylianakis *et al.* 2007), over environmental gradients Lurgi *et al.* 2012 and in time (Trøjelsgaard *et al.*, 2015) is accumulating rapidly. It is not clear, however, to what extent the turnover of network structure is driven by a systematic change in species composition or of pairwise interactions (Poisot *et al.*, 2012, 2015b). Our model comparison of host-parasite interactions revealed that most of the turnover is driven by species-specific responses to the environment, impacting species richness, and that co-occurrence was mostly neutral. Further, the occurrence of interactions among host and parasite is highly stochastic even when both are present, and not predictable by the variables considered by us. We know that interactions vary with the environment in other systems, for instance, herbivory (Shurin *et al.*, 2012) and predation (McKinnon *et al.*, 2010; Legagneux *et al.*, 2014) are often found to increase with temperature, resulting in spatial variation of trophic cascades (Gray *et al.*, 2015). What remains unclear, however, is the extent to which such variation is driven by a turnover of species composition along gradients, or a turnover of the interactions. Here we found that interactions vary substantially but non-predictably along the annual temperature and the precipitation gradient. Clearly, the lack of detectable signal may be due to our choice of covariates. Indeed, a previous study on a similar system identified habitat characteristics as the primary drivers of interactions (Nyman *et al.*, 2015). New investigations with other systems will thus be required to challenge this result. Under all circumstances, documenting the relationship between the environment and the occurrence of interactions at continental scales is critical for understanding how large-scale variation of trophic regulation influences community dynamics and ecosystem functioning (Harfoot *et al.*, 2014).

We restricted our framework to the effect of co-occurrence on ecological interactions, neglecting the inverse of the problem. We did not investigate in depth the drivers of co-occurrence and simply took it for granted from the data. Co-occurrence was indeed many times significantly different from the expectation of independent species distributions. It thus begs the question of whether, once environmental effects on species-specific distribution have been accounted for, interactions come with significant effects on co-occurrence? We could rephrase this problem by asking whether the fundamental niche differs from the realized niche, and how this applies to our framework. For example, we have considered above simply the co-occurrence probability, *P*(*X_iy_*, *X_jy_*|*E_y_*), which could be expanded as *P*(*X_iy_*|*X_jy_*, *E_y_*)*P*(*X_jy_*|*E_y_*). After some re-arrangement of Eq. 2, the marginal occurrence probability, *P*(*X_jy_*|*E_y_*), could be considered as a species distribution model taking into account the interaction between these species. This derivation would however critically depend on a strong *a priori* expectation of the conditional probability of observing a species given the distribution of the other species. This assumption seems reasonable for some situations, such as a parasitoid species that requires a host to develop. On the other hand, we found that the strength of this association is often rather weak if not neutral (for instance, with the example pair analyzed at Table 1). The lack of an association could simply arise when the parasitoid is generalist enough that it is not obligated to track the distribution of any single/given host (Cazelles *et al.*, 2015).

At present, there is only indirect support for the hypothesis that interacting species are conditionally distributed but this possibility should be the topic of more specific hypothesis testing. The impact of ecological interactions on the distribution of co-occurrence has been the topic of many publications since Diamond 1975 seminal study on competition and “checkerboard” distribution, but pairwise approaches have only recently received attention (Veech, 2013). Whether two interacting species are more closely associated in space remains unclear, since most approaches based on null models consider community-level metrics (e.g. Gotelli 2000), such as the C-score, thereby making it hard to evaluate if specific interactions do indeed affect co-occurrence. The expansion of the framework we describe to account for the difference between the realized and the fundamental niche will therefore require further investigation of the impact of interactions on co-occurrence.

Ecological networks are known to be extremely sparse, *i.e.* they have far more absences than presences of interactions. Absences of interactions, however, can come from different sources. The fact that unequal sampling at the local scale can affect our understanding of network structure is well documented (Martinez *et al.*, 1999). In a spatial context, however, some interactions may be undocumented simply because the species involved have never been observed to co-occur. Although these cases are reported as a lack of interactions, in actuality we cannot make any reliable inference from them: since the species have never been observed together, it remains possible that they would interact if they did. A fundamentally different category of absences of interactions are then those reported after multiple observations of species co-occurence. Thus, to gain confidence that the probability of an interaction is low, extensive sampling (that is, several records of co-occurence) is needed. Generally, our confidence that the interaction is indeed impossible will increase with the number of observations of the species pair. Seeing that this is essentially a Bernoulli process (the probability that the species will interact given their presence), the breadth of the confidence interval is expected to saturate after a fixed number of observations, which can be set as a threshold above which a species pair has finally been observed ‘often enough”. This will allow us to deal with both confirmed absences of interactions and mere absence of evidence.

## Conclusion

Our representation of spatial variation of community structure presents a new approach for the study of the biogeography of ecological networks. We see the following key challenges and opportunities ahead in this exciting area of research:

1. **New generation of network data.** Investigating spatial variation of network structure will require high quality and highly replicated network data. We have investigated one the most comprehensive spatial network datasets we are aware of and nonetheless found immense gaps of knowledge in its resolution. Species richness accumulates much faster than observations of ecological interactions (Poisot *et al.*, 2012). Each pair of species must be observed several times in order to obtain reliable estimates of their interaction probability.
2. **Estimation of the reliability of interactions.** We need quantitative tools to estimate the confidence intervals around inferred interaction probabilities, as well as estimators? of the frequency of false absences. Bayesian methods are promising to that end because we could use information on the target species (e.g. if they are known as specialists or generalists) to provide prior estimates of the interaction probability.
3. **From interaction probabilities to a distribution of network properties.** Metrics are available to analyze the structure of probabilistic networks (Poisot *et al.*, 2016a). These metrics are useful as first approximation, but they assume independence among interactions. This might not be the case in nature because of the role of cooccurrence and shared environmental requirements. We also need to better understand the distribution of network properties arising from probabilistic interactions.
4. **Investigation of the environmental-dependence of ecological interactions.** There is evidence that interactions can vary in space, but this problem has not been investigated in a systematic fashion. The paucity of currently available data precludes an extensive analysis of this question at present.
5. **Effects of ecological interactions on co-occurrence.** We have intentionally omitted the feedback of ecological interactions on co-occurrence in this framework. As abundance can impact the occurrence of interactions, and conversely since interactions impact abundance (Canard *et al.*, 2014), we could reasonably expect that interactions will also influence co-occurrence. Theory in this regard does exist for simple three-species modules (Cazelles *et al.*, 2015), but its extension to entire co-occurrence networks will prove critical in the future, especially given the interest in using co-occurrence to infer ecological interactions (Morales-Castilla *et al.*, 2015; Morueta-Holme *et al.*, 2016).

## Acknowledgements

This is a contribution to the working groups *Continental-scale variation of ecological networks* supported by the Canadian Institute for Ecology and Evolution), and the *Next Generation Data, Models, and Theory Working Group*, supported by the Santa Fe Institute, the Betsy and Jesse Fink Foundation, the ASU-SFI Center for Biosocial Complex Systems, and NSF Grant PHY-1240192. DG also acknowledge financial support from NSERC-Discovery grant program and Canada Research Chair program.

## References

Albouy,C., Velez,L., Coll,M., Colloca,F., Le Loc’h,F., Mouillot,D. & Gravel,D. (2014) From projected species distribution to food-web structure under climate change. Global Change Biology 20, 730–741.

Allesina,S., Alonso,D. & Pascual,M. (2008) A general model for food web structure. Science 320, 658–61.

Anderson,M.J., Crist,T.O., Chase,J.M., Vellend,M., Inouye,B.D., Freestone,A.L., Sanders,N.J., Cornell,H.V., Comita,L.S., Davies,K.F., Harrison,S.P., Kraft,N.J.B., Stegen,J.C. & Swenson,N.G. (2011) Navigating the multiple meanings of beta-diversity: a roadmap for the practicing ecologist. Ecology Letters 14, 19–28.

Argus,G. (1997) Infrageneric classification of Salix (Salicaceae) in the New World. Systematic Botany Monographs 52, 1–121.

Baiser,B., Gotelli,N.J., Buckley,H.L., Miller,T.E. & Ellison,A.M. (2012) Geographic variation in network structure of a nearctic aquatic food web. Global Ecology and Biogeography 21, 579–591.

Bartomeus,I., Gravel,D., Tylianakis,J., Aizen,M., Dickie,I. & Bernard-Verdier,M. (2016) A common framework for identifying linkage rules across different types of interactions. Functional Ecology pp. n/a–n/a.

Bascompte,J. & Jordano,P. (2007) Plant-Animal Mutualistic Networks: The Architecture of Biodiversity. Annual Review of Ecology, Evolution, and Systematics 38, 567–593.

Baselga,A. (2010) Partitioning the turnover and nestedness components of beta diversity. Global Ecology and Biogeography 19, 134–143.

Bersier,L.F., Banašek-Richter,C. & Cattin,M.F. (2002) Quantitative descriptors of food-web matrices. Ecology 83, 2394–2407.

Blonder,B., Lamanna,C., Violle,C. & Enquist,B.J. (2014) The n-dimensional hypervolume. Global Ecology and Biogeography 23, 595–609.

Boulangeat,I., Gravel,D. & Thuiller,W. (2012) Accounting for dispersal and biotic interactions to disentangle the drivers of species distributions and their abundances. Ecology Letters 15, 584–593.

Canard,E., Mouquet,N., Mouillot,D., Stanko,M., Miklisova,D. & Gravel,D. (2014) Empirical evaluation of neutral interactions in host-parasite networks. The American naturalist 183, 468–79.

Cazelles,K., Araújo,M.B., Mouquet,N. & Gravel,D. (2016) A theory for species cooccurrence in interaction networks. Theoretical Ecology 9, 39–48.

Cazelles,K., Mouquet,N., Mouillot,D. & Gravel,D. (2015) On the integration of biotic interaction and environmental constraints at the biogeographical scale. Ecography pp. n/a–n/a.

Chase,J. & Leibold,M. (2003) Ecological niches. Chicago University Press, Chicago.

Chase,J.M. (2007) Drought mediates the importance of stochastic community assembly. Proceedings of the National Academy of Sciences of the United States of America 104, 17430–4.

Connor,E. & Simberloff,D. (1979) The assembly of species communities: chance or competition? Ecology 60, 1132–1140.

Cronk,Q., Ruzzier,E., Belyaeva,I. & Percy,D. (2015) Salix transect of Europe: latitudinal patterns in willow diversity from Greece to arctic Norway. Biodiversity Data Journal 3.

Diamond,J. (1975) Assembly of species communities. Ecology and evolution of communities (eds. M. Cody & J. Diamond), pp. 342–444, Harvard University Press, Cambridge.

Dunne,J.A. (2006) The network structure of food webs. Ecological networks: Linking structure and dynamics (eds. M. Pascual & J.A. Dunne), pp. 27–86, Oxford University Press, Oxford.

Eklöf,A., Jacob,U., Kopp,J., Bosch,J., Castro-Urgal,R., Chacoff,N.P., Dalsgaard,B., de Sassi,C., Galetti,M., Guimarães,P.R., Lomáscolo,S.B., Martín González,A.M., Pizo,M.A., Rader,R., Rodrigo,A., Tylianakis,J.M., Vázquez,D.P. & Allesina,S. (2013) The dimensionality of ecological networks. Ecology letters 16, 577–583.

Gibert,J.P. & DeLong,J.P. (2014) Temperature alters food web body-size structure. Biology Letters 10.

Gleason,H. (1926) The individualistic concept of the plant association. Bulletin of the Torrey Botanical Club 53, 7–26.

Gotelli,N.J. (2000) Null Model Analysis of Species Co-Occurrence Patterns. Ecology 81, 2606.

Götzenberger,L., de Bello,F., Bråthen,K.A., Davison,J., Dubuis,A., Guisan,A., Lepå,J., Lindborg,R., Moora,M., Pårtel,M., Pellissier,L., Pottier,J., Vittoz,P., Zobel,K. & Zobel,M. (2012) Ecological assembly rules in plant communities-approaches, patterns and prospects. Biological reviews of the Cambridge Philosophical Society 87, 111–27.

Gravel,D., Massol,F., Canard,E., Mouillot,D. & Mouquet,N. (2011) Trophic theory of island biogeography. Ecology letters 14, 1010–6.

Gravel,D., Poisot,T., Albouy,C., Velez,L. & Mouillot,D. (2013) Inferring food web structure from predator-prey body size relationships. Methods in Ecology and Evolution 4, 1083–1090.

Gray,S.M., Poisot,T., Harvey,E., Mouquet,N., Miller,T.E. & Gravel,D. (2015) Temperature and trophic structure are driving microbial productivity along a biogeographical gradient. Ecography pp. n/a–n/a.

Hanski,I. (1999) Metapopulation Ecology. Oxford University Press, Oxford.

Hardin,G. (1960) The competitive exclusion principle. Science 131, 1292–1297.

Harfoot,M.B.J., Newbold,T., Tittensor,D.P., Emmott,S., Hutton,J., Lyutsarev,V., Smith,M.J., Scharlemann,J.P.W. & Purves,D.W. (2014) Emergent global patterns of ecosystem structure and function from a mechanistic general ecosystem model. PLoS Biol 12, 1–24.

Havens,K. (1992) Scale and structure in natural food webs. Science 257, 1107–9.

Hijmans,R.J. (2015) raster: Geographic Data Analysis and Modeling. R package version 2.5–2.

Holt,R.D. (2009) Bringing the Hutchinsonian niche into the 21st century: Ecological and evolutionary perspectives. Proceedings of the National Academy of Sciences 106, 19659–19665.

Holt,R.D. & Lawton,J.H. (1993) Apparent competition and enemy-free space in insect host-parasitoid communities. The American Naturalist 142, 623–645, pMID: 19425963.

Hutchinson,G. (1957) Concluding remarks. Cold Spring Harbour Symposium 22, 415–427.

Ings,T.C., Montoya,J.M., Bascompte,J., Blüthgen,N., Brown,L., Dormann,C.F., Edwards,F., Figueroa,D., Jacob,U., Jones,J.I., Lauridsen,R.B., Ledger,M.E., Lewis,H.M., Olesen,J.M., van Veen,F.J.F., Warren,P.H. & Woodward,G. (2009) Ecological networks-beyond food webs. The Journal of animal ecology 78, 253–69.

Kéfi,S., Berlow,E.L., Wieters,E.a., Navarrete,S.a., Petchey,O.L., Wood,S.a., Boit,A., Joppa,L.N., Lafferty,K.D., Williams,R.J., Martinez,N.D., Menge,B.a., Blanchette,C.a., Iles,A.C. & Brose,U. (2012) More than a meal…integrating non-feeding interactions into food webs. Ecology letters pp. 291–300.

Kissling,W.D., Dormann,C.F., Groeneveld,J., Hickler,T., Kuhn,I., McInerny,G.J., Montoya,J.M., Romermann,C., Schiffers,K., Schurr,F.M., Singer,A., Svenning,J.C., Zimmermann,N.E. & O’Hara,R.B. (2012) Towards novel approaches to modelling biotic interactions in multispecies assemblages at large spatial extents. Journal of Biogeography 39, 2163–2178.

Kopelke,J.P. (1985) Über die Biologie und Parasiten der gallenbildenden Blattwespe-naften Pontania dolichura (Thoms, 1871), P. vesicator (Bremi 1849) und P. viminalis (L. 1758) (Hymenoptera: Tenthredinidae). Faunistisch-Okologische Mitteilungen 5, 331–344.

Kopelke,J.P. (1999) Gallenerzeugende Blattwespen Europas-Taxonomische Grundlagen, Biologie und Ökologie (Tenthredinidae: Nematinae: Euura, Phyllocolpa, Pontania). Courier Forschungsinstitut Senckenberg 212, 1–183.

Kopelke,J.P. (2003) Natural enemies of gall-forming sawflies on willows (Salix spp.). Entomologia Generalis 26, 277–312.

Kouki,J. (1999) Latitudinal gradients in species richness in northern areas: some exceptional patterns. Ecological Bulletins 47, 30–37.

Kouki,J., Niemelä,P. & Viitasaari,M. (1994) Reversed latitudinal gradient in species richness of sawflies (Hymenoptera, Symphyta). Annales Zoologici Fennici 31, 83–88.

Krebs,C. (2009) Ecology: The Experimental Analysis of Distribution and Abundance. 6th ed. Pearson, San Francisco.

Lalibertä,E. & Tylianakis,J.M. (2010) Deforestation homogenizes tropical parasitoid-host networks. Ecology 91, 1740–1747.

Legagneux,P., Gauthier,G., Lecomte,N., Schmidt,N.M., Reid,D., Cadieux,M.C., Berteaux,D., Bêty,J., Krebs,C.J., Ims,R.a., Yoccoz,N.G., Morrison,R.I.G., Leroux,S.J., Loreau,M. & Gravel,D. (2014) Arctic ecosystem structure and functioning shaped by climate and herbivore body size. Nature Climate Change E2168, 1–5.

Legendre,P., Borcard,D. & Peres-Neto,P.R. (2005) Analyzing beta diversity: partitioning the spatial variation of composition data. Ecological Monographs 75, 435–450.

Legendre,P. & De Caceres,M. (2013) Beta diversity as the variance of community data: dissimilarity coefficients and partitioning. Ecology Letters 16, 951–963.

Leibold,M., Holyoak,M., Mouquet,N., Amarasekare,P., Chase,J.M., Hoopes,M., Holt,R.D., Shurin,J.B., Law,R., Tilman,D., Loreau,M. & Gonzalez,A. (2004) The metacommunity concept: a framework for multi-scale community ecology. Ecology Letters 7, 601–613.

Leppänen,S., Malm,T., Värri,K. & Nyman,T. (2014) A comparative analysis of genetic differentiation across six shared willow host species in leaf-and bud-galling sawflies. PLoS ONE 9, e116286.

Lurgi,M., Läpez,B.C. & Montoya,J.M. (2012) Novel communities from climate change. Philosophical transactions of the Royal Society of London. Series B, Biological sciences 367, 2913–22.

Martinez,N.D., Hawkins,B.a., Dawah,H.A. & Feifarek,B.P. (1999) Effects of Sampling Effort on Characterization of Food-Web Structure. Ecology 80, 1044–1055.

McInerny,G.J. & Etienne,R.S. (2012a) Ditch the niche-is the niche a useful concept in ecology or species distribution modelling? Journal of Biogeography 39, 2096–2102.

McInerny,G.J. & Etienne,R.S. (2012b) Stitch the niche-a practical philosophy and visual schematic for the niche concept. Journal of Biogeography 39, 2103–2111.

McKinnon,L., Smith,P.A., Nol,E., Martin,J.L., Doyle,F.I., Abraham,K.F., Gilchrist,H.G., Morrison,R.I.G. & Bety,J. (2010) Lower predation risk for migratory birds at high latitudes. Science 327, 326–327.

Morales-Castilla,I., Matias,M.G., Gravel,D. & Araújo,M.B. (2015) Inferring biotic interactions from proxies. Trends in Ecology & Evolution 30, 347–356.

Morueta-Holme,N., Blonder,B., Sandel,B., McGill,B.J., Peet,R.K., Ott,J.E., Violle,C., Enquist,B.J., Jørgensen,P.M. & Svenning,J.C. (2016) A network approach for inferring species associations from co-occurrence data. Ecography pp. n/a–n/a.

Nyman,T., Bokma,F. & Kopelke,J.P. (2007) Reciprocal diversification in a complex plant-herbivore-parasitoid food web. BMC Biology 5, 49.

Nyman,T. & Julkunen-Tiitto,R. (2000) Manipulation of the phenolic chemistry of willows by gall-inducing sawflies. Proceedings of the National Academy of Sciences 97, 13184–13187.

Nyman,T., Leppänen,S.A., Värkonyi,G., Shaw,M.R., Koivisto,R., Barstad,T.E., Vikberg,V. & Roininen,H. (2015) Determinants of parasitoid communities of willow-galling sawflies: habitat overrides physiology, host plant and space. Molecular Ecology 24, 5059–5074.

Nyman,T., Vikberg,V., Smith,D. & Boevä J-L (2010) How common is ecological speci-ation in plant-feeding insects? A “Higher” Nematinae perspective. BMC Evolutionary Biology 10, e266.

Pellissier,L., Rohr,R., Ndiribe,C., Pradervand,J.N., Salamin,N., Guisan,A. & Wisz,M. (2013) Combining food web and species distribution models for improved community projections. Ecology and Evolution 3, 4572–4583.

Petchey,O.L., Brose,U. & Rall,B.C. (2010) Predicting the effects of temperature on food web connectance. Philosophical transactions of the Royal Society of London. Series B, Biological sciences 365, 2081–91.

Peters,R. (1991) A critique for ecology. Cambridge University Press, Cambridge.

Phillips,S.J., Anderson,R.P. & Schapire,R.E. (2006) Maximum entropy modeling of species geographic distributions. Ecological Modelling 190, 231–259.

Pillai,P., Gonzalez,A. & Loreau,M. (2011) Metacommunity theory explains the emergence of food web complexity. Proceedings of the National Academy of Sciences of the United States of America 108, 19293–19298.

Poisot,T., Baiser,B., Dunne,J.A., Kéfi,S., Massol,F., Mouquet,N., Romanuk,T.N., Stouffer,D.B., Wood,S.A. & Gravel,D. (2015a) mangal-making ecological network analysis simple. Ecography 37.

Poisot,T., Canard,E., Mouillot,D., Mouquet,N. & Gravel,D. (2012) The dissimilarity of species interaction networks. Ecology letters XX, XX–XX.

Poisot,T., Cirtwill,A.R., Cazelles,K., Gravel,D., Fortin,M.J. & Stouffer,D.B. (2016a) The structure of probabilistic networks. Methods in Ecology and Evolution 7, 303–312.

Poisot,T. & Gravel,D. (2014) When is an ecological network complex? connectance drives degree distribution and emerging network properties. PeerJ 2, e251.

Poisot,T., Gravel,D., Leroux,S., Wood,S.A., Fortin,M.J., Baiser,B., Cirtwill,A.R., Araujo,M.B. & Stouffer,D.B. (2016b) Synthetic datasets and community tools for the rapid testing of ecological hypotheses. Ecography 39, 402–408.

Poisot,T., Stouffer,D.B. & Gravel,D. (2015b) Beyond species: why ecological interactions vary through space and time. Oikos 124, 243–251.

Pollock,L.J., Tingley,R., Morris,W.K., Golding,N., O’Hara,R.B., Parris,K.M., Vesk,P.A. & McCarthy,M.A. (2014) Understanding co-occurrence by modelling species simultaneously with a joint species distribution model (jsdm). Methods in Ecology and Evolution 5, 397–406.

Pulliam,H.R. (2000) On the relationship between niche and distribution. Ecology Letters 3, 349–361.

Roininen,H., Nyman,T. & Zinovjev,A. (2005) Biology, ecology, and evolution of gall-inducing sawflies (Hymenoptera: Tenthredinidae and Xyelidae). Biology, Ecology, and Evolution of Gall-Inducing Arthropods (eds. A.Raman, C.W. Schaefer & T.M. Withers), pp. 467–494, Science Publishers Inc.,

Scheiner,S.M. & Willig,M.R. (2007) A general theory of ecology. Vital And Health Statistics. Series 20 Data From The National Vitalstatistics System Vital Health Stat 20 Data Natl Vital Sta.

Shurin,J.B., Clasen,J.L., Greig,H.S., Kratina,P. & Thompson,P.L. (2012) Warming shifts top-down and bottom-up control of pond food web structure and function. Philosophical Transactions of the Royal Society of London B: Biological Sciences 367, 3008–3017.

Silvertown,J. (2004) Plant coexistence and the niche. Trends in Ecology and Evolution 19, 605–611.

Skvortsov,A.K. (1999) Willows of Russia and adjacent countries. Taxonomical and geographical revision. Univ. Joensuu Fac. Math. Natl. Sci. Rep. Ser 39, 1–307.

Soberón,J. (2007) Grinnellian and eltonian niches and geographic distributions of species. Ecology Letters 10, 1115–1123.

Stouffer,D.B., Camacho,J., Jiang,W. & Amaral,L.a.N. (2007) Evidence for the existence of a robust pattern of prey selection in food webs. Proceedings of the Royal Society B: Biological Sciences 274, 1931–40.

Thuiller,W. (2003) Biomod-optimizing predictions of species distributions and projecting potential future shifts under global change. Global Change Biology 9, 1353–1362.

Tilman,D. (1982) Resource competition and community structure. Princeton University Press, Princeton.

Townsend,P.A., Soberón,S., Pearson,R.G., Anderson,R.P., Martínez-Meyer,E., Miguel,N. & Araújo,M.B. (2011) Ecological Niches and Geographic Distributions. Princeton University Press, Princeton.

Trøjelsgaard,K., Jordano,P., Carstensen,D.W. & Olesen,J.M. (2015) Geographical variation in mutualistic networks: similarity, turnover and partner fidelity. Proceedings of the Royal Society of London B: Biological Sciences 282, 20142925.

Tylianakis,J.M., Tscharntke,T. & Lewis,O.T. (2007) Habitat modification alters the structure of tropical host-parasitoid food webs. Nature 445, 202–205.

Veech,J.A. (2013) A probabilistic model for analysing species co-occurrence. Global Ecology and Biogeography 22, 252–260.

Veijalainen,A., Súúksjarvi,I., Erwin,T., Gúmez,I. & Longino,J. (2013) Subfamily composition of Ichneumonidae (Hymenoptera) from western Amazonia: insights into diversity of tropical parasitoid wasps. Insect Conservation and Diversity 6, 28–37.

Vellend,M., Srivastava,D.S., Anderson,K.M., Brown,C.D., Jankowski,J.E., Kleyn-hans,E.J., Kraft,N.J.B., Letaw,A.D., Macdonald,A.A.M., Maclean,J.E., Myers-Smith,I.H., Norris,A.R. & Xue,X. (2014) Assessing the relative importance of neutral stochasticity in ecological communities. Oikos 123, 1420–1430.

Williams,R. & Martinez,N. (2000) Simple rules yield complex food webs. Nature 404, 180–183.

Williams,R.J., Anandanadesan,A. & Purves,D. (2010) The probabilistic niche model reveals the niche structure and role of body size in a complex food web. PloS One 5, e12092.

Wisz,M.S., Pottier,J., Kissling,W.D., Pellissier,L., Lenoir,J., Damgaard,C.F., Dor-mann,C.F., Forchhammer,M.C., Grytnes,J.A., Guisan,A., Heikkinen,R.K., Høye,T.T., Kuhn,I., Luoto,M., Maiorano,L., Nilsson,M.C., Normand,S., Ockinger,E., Schmidt,N.M., Termansen,M., Timmermann,A., Wardle,D.a., Aastrup,P. & Sven-ning,J.C. (2012) The role of biotic interactions in shaping distributions and realised assemblages of species: implications for species distribution modelling. Biological Reviews.

Wood,S.A., Russell,R., Hanson,D., Williams,R.J. & Dunne,J.A. (2015) Effects of spatial scale of sampling on food web structure. Ecology and Evolution 5, 3769–3782.

Woodward,G., Perkins,D.M. & Brown,L.E. (2010) Climate change and freshwater ecosystems: impacts across multiple levels of organization. Philosophical transactions of the Royal Society of London. Series B, Biological sciences 365, 2093–106.

Wu,J., Nyman,T., Wang,D.C., Argus,G., Yang,Y.P. & Chen,J.H. (2015) Phylogeny of Salix subgenus Salix s.l. (Salicaceae): delimitation, biogeography, and reticulate evolution. BMC Evolutionary Biology 15, e31.

